# A phenotypic screen of Marfan syndrome iPSC-derived vascular smooth muscle cells uncovers GSK3β as a new target

**DOI:** 10.1101/2022.04.11.487841

**Authors:** Hongorzul Davaapil, Madeline McNamara, Alessandra Granata, Robyn G.C. Macrae, Mei Hirano, Martina Fitzek, Jose Antonio Aragon-Martin, Anne Child, David M. Smith, Sanjay Sinha

## Abstract

Marfan syndrome (MFS) is a rare connective tissue disorder caused by mutations in *FBN1*. Patients with MFS notably suffer from aortic aneurysm and dissection. Despite considerable effort, animal models have proven to be poorly predictive for therapeutic intervention in human aortic disease. Using a “humanised” model system may be more appropriate in identifying new therapeutic targets. Patient-derived induced pluripotent stem cells can be differentiated into vascular smooth muscle cells (VSMCs) and recapitulate major features of MFS. We have screened 1,022 small molecules in our *in vitro* model, exploiting the highly-proteolytic nature of MFS-VSMCs, and identified 36 effective compounds. Further analysis identified GSK3β as a recurring target in the compound screen. GSK3β inhibition/knockdown did not ameliorate the proliferation defect in MFS-VSMCs but improved MFS-VSMC apoptosis and proteolysis. To conclude, we have identified GSK3β as a novel target for MFS, forming the foundation for future work in MFS and other aortic diseases.

## Introduction

Marfan syndrome (MFS) is a rare genetic disorder resulting in multi-system abnormalities. It is caused by deleterious variants in the *FBN1* gene, a key extracellular matrix (ECM) protein in connective tissue (Dietz et al., 1992). The cardiovascular effects can be life-threatening, as patients can develop thoracic aortic aneurysm and dissection (TAAD), particularly at the aortic root and arch. It is currently thought that the majority of aortic disease is propagated through vascular smooth muscle cells (VSMCs), although there is also evidence of endothelial dysfunction (Chung et al., 2007; Galatioto et al., 2018; Oller et al., 2017). In addition, there is heterogeneity in the embryonic origin of VSMCs present in aorta, which itself has been hypothesised to contribute to disease progression (Majesky, 2007).

The current treatment options for patients with MFS are limited to prescription of anti-hypertensives, surgical replacement or external support (Pepper et al., 2020) of the dilated aortic root – a major procedure with significant risk of morbidity and mortality. The use of the angiotensin II receptor blocker (ARB) losartan in a mouse model of MFS was highly effective in limiting aortic disease progression (Habashi et al., 2006). Unfortunately, following up from this work, numerous clinical trials have concluded that losartan was either not successful or had only modest effects in reducing aortic diameter or improving clinical end-points in patients (Groenink et al., 2013; Lacro et al., 2014; Milleron et al., 2015; Teixido-Tura et al., 2018). These disappointing results could be attributed to a variety of factors, including insufficient safe dosage (Mullen et al., 2020), fundamental differences between the species and varying genetic backgrounds. There is therefore a need for alternative approaches to identify novel and effective treatment options for MFS.

Induced pluripotent stem cells (iPSCs) can be used to generate any somatic cell type, including lineage-specific VSMCs. We have developed protocols to generate lateral plate mesoderm, neural crest (NC) and paraxial mesoderm-derived VSMCs, which correspond to the aortic root, ascending aorta and descending aorta respectively (Cheung et al., 2012, 2014). Using this lineage-specific approach, an iPSC-based model of MFS *in vitro* has been developed (Granata et al., 2017). There, the main features of the aortic phenotype in VSMCs were recapitulated in NC-VSMCs; notably, abnormal ECM deposition, increased matrix metalloproteinase (MMP) expression and activity, apoptosis and abnormal response to mechanical stretch. We identified p38 as a candidate for mediating the MFS phenotype, as p38 inhibition partially rescued the phenotype *in vitro* (Granata et al., 2017).

Here, we describe a medium-throughput, unbiased small molecule (SM) screen to identify novel disease mechanisms and therapeutic targets using iPSC technology. In collaboration with AstraZeneca, we have screened 1,022 small molecules on MFS NC-VSMCs and identified a subset which were found to reduce MMP activity. In particular, we identified that GSK3β SM inhibitors (SMIs) and genetic knockdown improved cellular function, where MFS-VSMCs were less proteolytic and showing reduced apoptosis. In addition, we treated three additional MFS patient lines with a GSK3β inhibitor and obtained a consistent outcome, suggesting that this may be a common cellular defect amongst different MFS patient lines. This work highlights a screening strategy which could be used widely to screen additional SMIs and/or applied to models of other aortic diseases.

## Results

### Screen of 1,022 SMs

SMs are low molecular weight compounds, typically of around 500 Da in size, which can modulate protein binding and activity (Khera and Rajput, 2017; Zhong et al., 2021). Because of their low molecular weight, they are able to penetrate cells more easily than macromolecular drugs, such as antibodies or other proteins. Here, we sought to screen a library of 1,022 SMs to identify any compounds which can ameliorate the disease phenotype of MFS VSMCs. The library of compounds used in this study was obtained from AstraZeneca’s Open Innovation Group. It is composed of 14,000 SMs in total and was recently used for SM screening in an iPSC model of non-alcoholic fatty liver disease (Parafati et al., 2020). These compounds are highly annotated and have information on pIC50 for primary and secondary targets; over 1,700 targets are covered in all. This library is composed of SMIs which target multiple proteins within a given signalling pathway, thereby increasing its capacity to uncover new pathways implicated in disease. As there is significant over-representation of some targets, the library of 14,000 SMIs was selectively narrowed down to 1,022 compounds in order to maintain a broad cohort of targets in a smaller number of compounds for this proof-of-concept work.

We designed this phenotypic screen around the highly-proteolytic nature of MFS VSMCs, which have elevated MMP expression and secretion (Cui et al., 2021; Granata et al., 2017; Ikonomidis et al., 2006). The patient line used for this study, unless specified, is the FBN1 C1242Y line which we have previously characterised (Granata et al., 2017). A fluorescence-quenched gelatine substrate would then be incubated with MMPs from the cell culture medium: cleavage of this substrate would result in a fluorescent signal which can be detected in a plate-reader (Figure 1B). MFS VSMCs were treated for 96 hours with 1μM SM, after which the supernatant was collected and analysed. Of the 1022 compounds tested at this concentration for 96 hours, 730 were found to be associated with some cell atrophy and/or detachment which made them poor candidates for proceeding to assay for MMP activity. Of the remaining 292 SMs, 36 were found to successfully reduce MMP activity down to levels comparable to the isogenic corrected control (Corr) VSMCs, or MFS VSMCs treated with losartan (Figure 1C & D).

**Figure 1.**
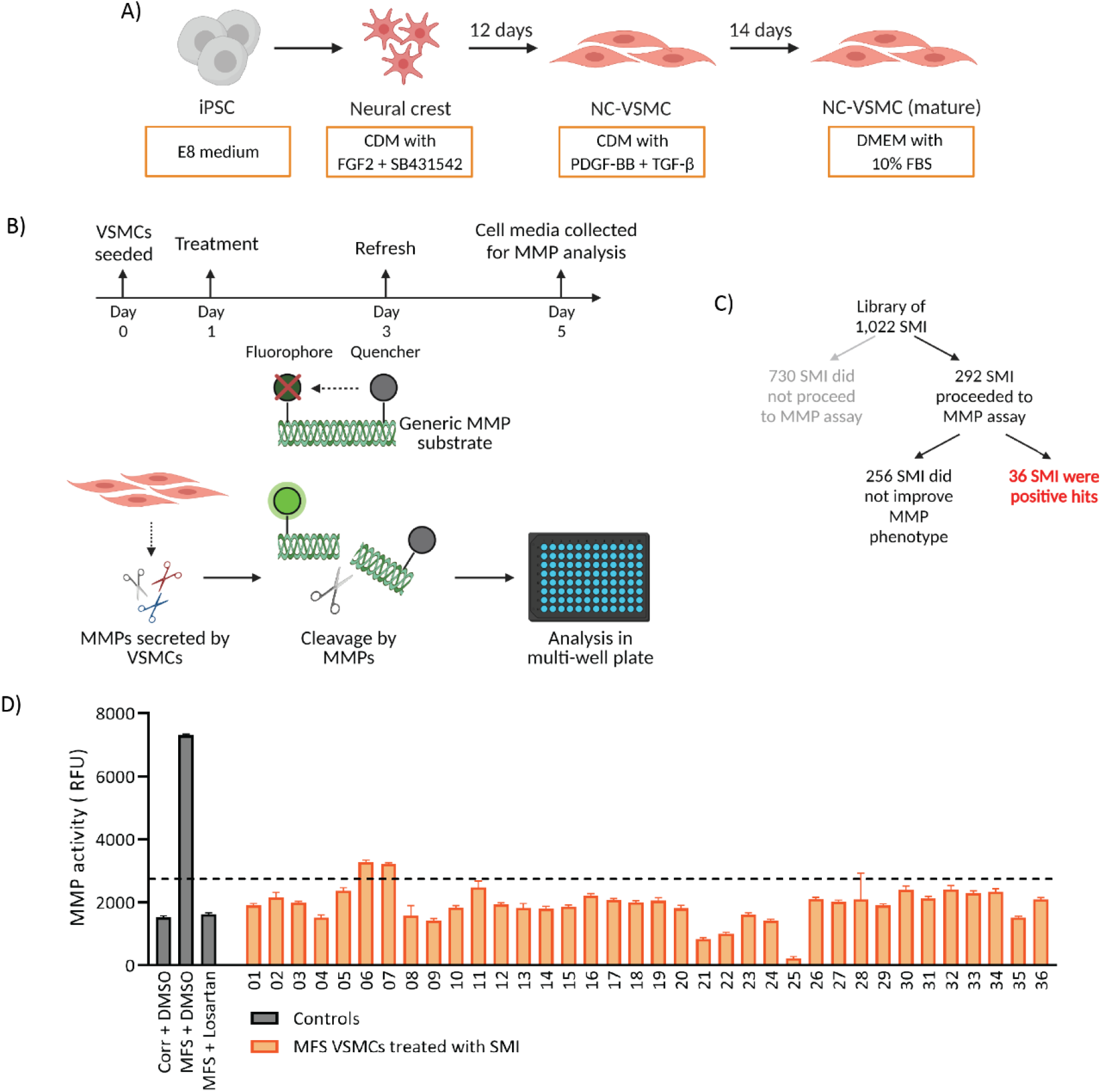
MMP activity-based drug screen. **A)** Overview of differentiation to NC-VSMCs from iPSCs. **B)** MFS VSMCs were treated with 1μM SMIs for 96 hours and cell supernatant collected. This supernatant contains secreted MMPs which, when incubated with the generic MMP substrate, leads to cleavage and subsequent fluorescent signal, which can be measured using a plate-reader. **C)** Of the 1,022 SMIs, the majority were not suitable for further assay at 1μM. **D)** Among the SMIs used in the screen, 36 were found to decrease MMP activity of MFS VSMCs. Corrected VSMCs and MFS VSMCs treated with losartan were used as controls to determine the threshold of sufficient MMP activity reduction. n=2. Data are represented as mean ± SEM.

### GSK3β is a recurring target among the positive hits

In order to identify a promising target worthy of further investigation, we analysed the annotated primary and secondary targets and their associated pIC50 values (Data 1). From the 36 SMIs, we identified 902 unique targets (Figure 2A). Since SMs were used at 1μM, pIC50 values below 6 are not informative – therefore, drug targets with a pIC50 < 6 were not included in our analysis, resulting in 538 unique targets.

**Figure 2.**
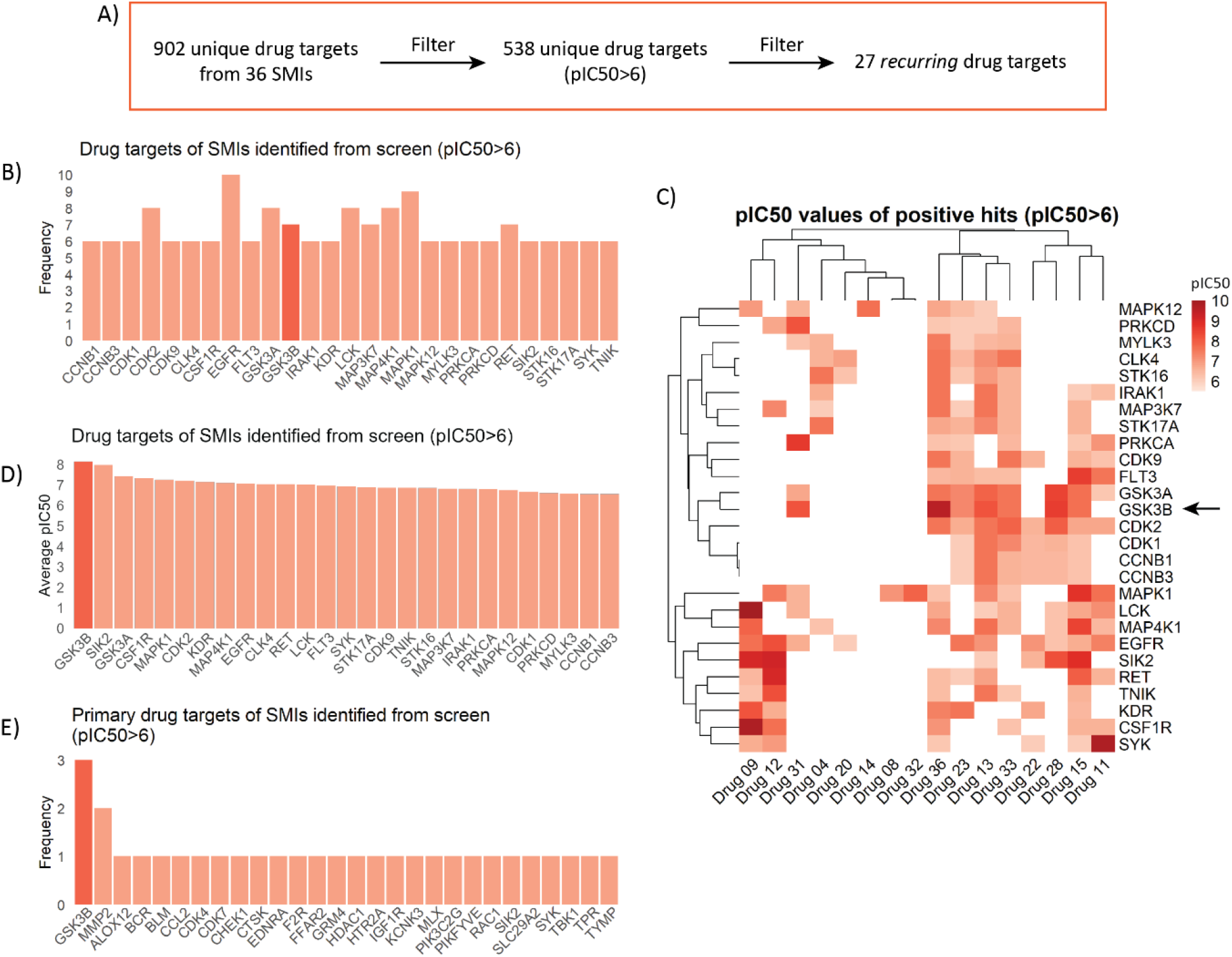
GSK3β is a recurring drug target among the positive hits from the drug screen. **A)** Outline of how drug targets were filtered. This was performed by removing drug targets with pIC50<6, and then overall frequency lower than 5. **B)** Frequency of drug targets suggests that GSK3β (red) is a recurring target amongst others. **C)** Heatmap of pIC50 values for all high-frequency drug targets indicates that GSK3β (arrow) is also a high-specificity drug target. **D)** Average pIC50 values for each drug target indicates that GSK3β (red) was the most specific target. **E)** Primary drug targets amongst the positive hits from the drug screen. GSK3β (red) is the most recurring primary target.

The majority of these targets were protein kinases, as indicated by GO term enrichment (Supp Figure 1A). Interestingly, we identified p38 MAPK inhibitors amongst our positive hits, along with GABA receptor inhibitors, both of which have been found to be effective in MFS by us and others (Supp Figure 1B) (Granata et al., 2017; Hansen et al., 2019). KEGG pathway enrichment analysis indicates that components of the MAPK signal transduction pathway are highly enriched, along with other potentially interesting pathways, such as those linked to focal adhesions (Supp Figure 1C).

Out of 538 unique targets, 27 targets were found to be present in 6 or more SMs (Figure 2B; Data 2). We identified that GSK3β is a recurrent and highly-specific target, as illustrated by the heatmap (Figure 2C; arrow) and average pIC50 values (Figure 2D; red). Amongst the negative hits from the SM screen, i.e. compounds which did not produce a beneficial effect in MFS VSMCs, GSK3β was not a top recurring target (Supp Figure 1D). In addition, we also found that GSK3β was the most prominent primary target amongst the SMIs (Figure 2E). Finally, correlation between positive and negative hits (Supp Figure 2) also demonstrated that GSK3β is a top contender for consideration, which is emphasised by using a more stringent pIC50 threshold (Supp Figure 2). This is particularly important as the pIC50 values from this dataset are derived from isolated enzyme activity assays – it is likely that the activity of compounds inside the cell would be lower. This reinforces our approach to use pIC50 as a cut-off, but also suggests that even higher stringency may also be informative. Taken together, we therefore decided to focus our validation on GSK3β.

### GSK3β expression in Corr vs MFS cells

We started by assessing the expression of GSK3β in untreated Corr and MFS VSMCs. At the mRNA level, GSK3β expression trended towards an increase (p=0.07) in MFS cells (Figure 3A). In contrast, total GSK3β expression in MFS VSMCs is decreased compared to the corrected control (Figure 3B). Interestingly, it seems that GSK3β activity is decreased too: there is increased phosphorylation at Ser9 (Figure 3B), which is an inhibitory post-translational modification (Cross et al., 1995). This is supported by the increased amount of β-catenin in MFS cells, indicative of increased signalling through the canonical Wnt pathway. This finding was unexpected, as it suggests that GSK3β activity may already be decreased in the MFS cells, yet GSK3β inhibition was identified from the drug screen as effective at reversing the MFS proteolytic phenotype.

**Figure 3.**
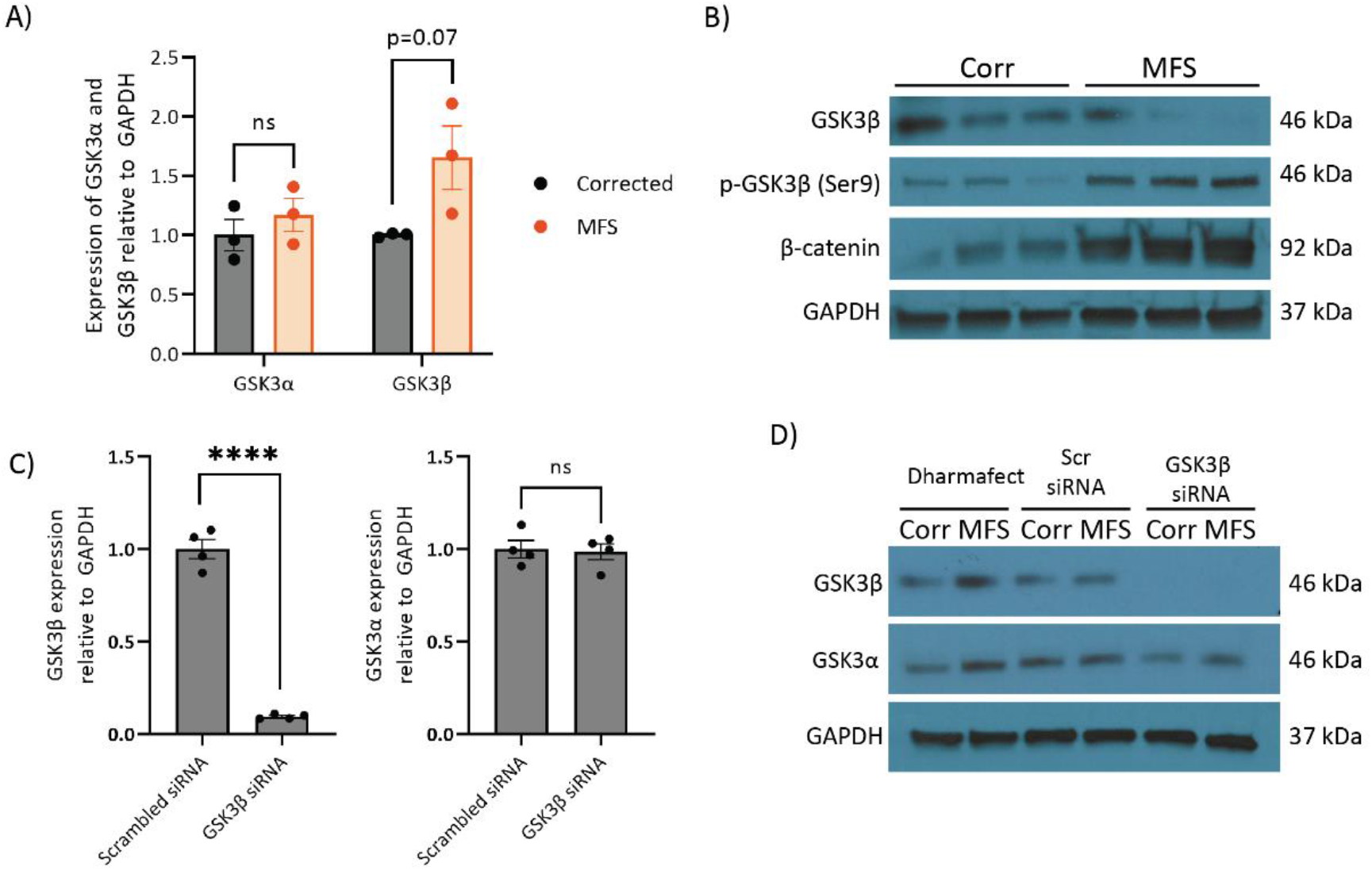
GSK3β expression in MFS VSMCs and its knockdown by siRNA. **A)** Expression of GSK3α and GSK3β mRNA in Corrected and MFS VSMCs. **B)** Expression of GSK3β, phospho-GSK3β (Ser9) and β-catenin protein in corrected and MFS VSMCs. GAPDH was used as a loading control. n=3 for both qPCR and western blotting. Compared to the scrambled control siRNA, siRNA against GSK3β was effective in knocking down its expression at both the mRNA **(C)** and protein **(D)** levels, without altering the expression of GSK3α. GAPDH used as the loading control. Corrected cells (n=4) used for qPCR analysis, and representative corrected and MFS cells used for western blotting. Data are represented as mean ± SEM.

Next, in order to help with validation of GSK3β as a target, we used siRNA to knockdown the expression of GSK3β. Since SMIs frequently have multiple secondary drug targets, we used a genetic system to also verify and validate the results of our SM screen. Furthermore, knockdown was used instead of CRISPR-mediated deletion as this would not abrogate expression entirely, mimicking the effects more closely of inhibition by SMIs. siRNA-mediated knockdown of GSK3β was successful at reducing the expression of both mRNA and protein (Figure 3C). We also confirmed that this strategy did not affect the expression of GSK3α (Figure 3D).

### Decreased GSK3β reduces MMP activity and expression

Next, we aimed to confirm the findings of the drug screen. In addition to siRNA-mediated knockdown, we decided to use six SMIs which target GSK3β: three inhibitors identified from the drug screen (6BIO, AZ1 and AZ2), along with three additional compounds (CHIRON, AZ3 and AZ4) for further validation (Table 1). In order to do this, we performed *in-situ* zymography, where we cultured cells on DQ-gelatin. Similar to the MMP substrate used for the initial screen, DQ-gelatin fluoresces when cleaved by MMPs, resulting in deposition of green fluorescence. After 96 hours of treatment, cells were imaged, and the data quantified in an automated and unbiased manner. Our findings indicate that while the MFS VSMCs treated with DMSO or scrambled siRNA exhibited high levels of DQ-gelatin degradation, cells treated with the GSK3β inhibitors (1μM) or siRNA showed less degradation, with levels similar to those of corrected control cells (Figure 4A, B & C). This successfully recapitulated decreased matrix degradation observed when cells were treated with doxycycline, losartan and p38 inhibitor losmapimod (Supp Figure 3). This finding was further supported by decreased expression of MMPs 2 and 9 upon treatment with GSK3β inhibitors (Figure 4D & E). These results therefore confirm that GSK3β inhibition is beneficial in decreasing the proteolytic nature of MFS VSMCs.

**Table 1.**
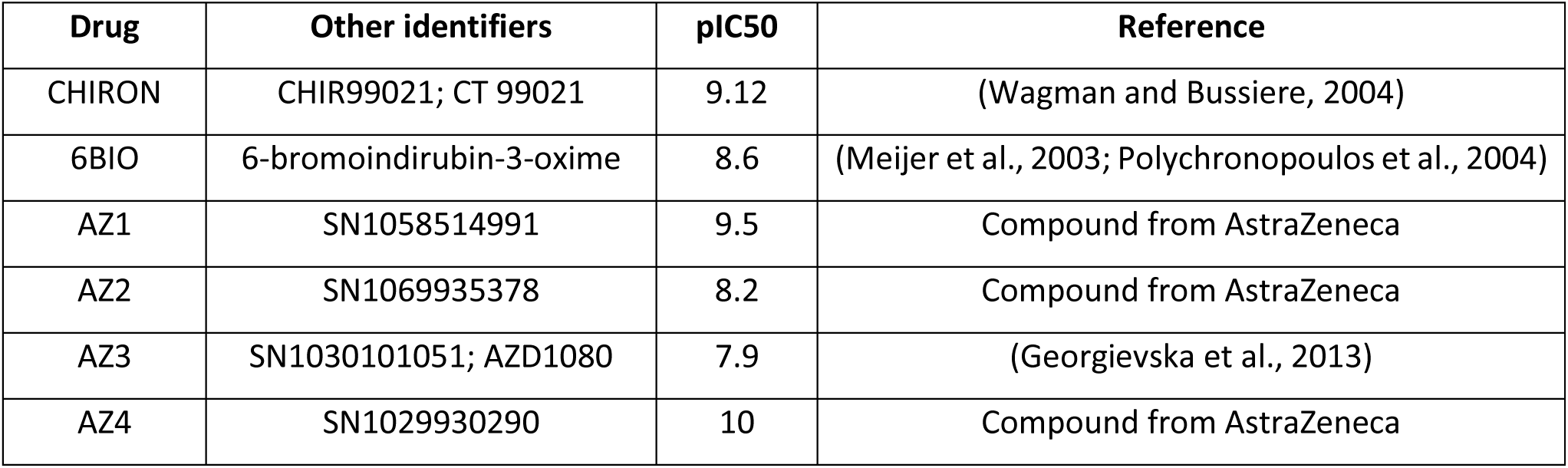
GSK3β SMI used and their pIC50 values for GSK3β.

**Figure 4.**
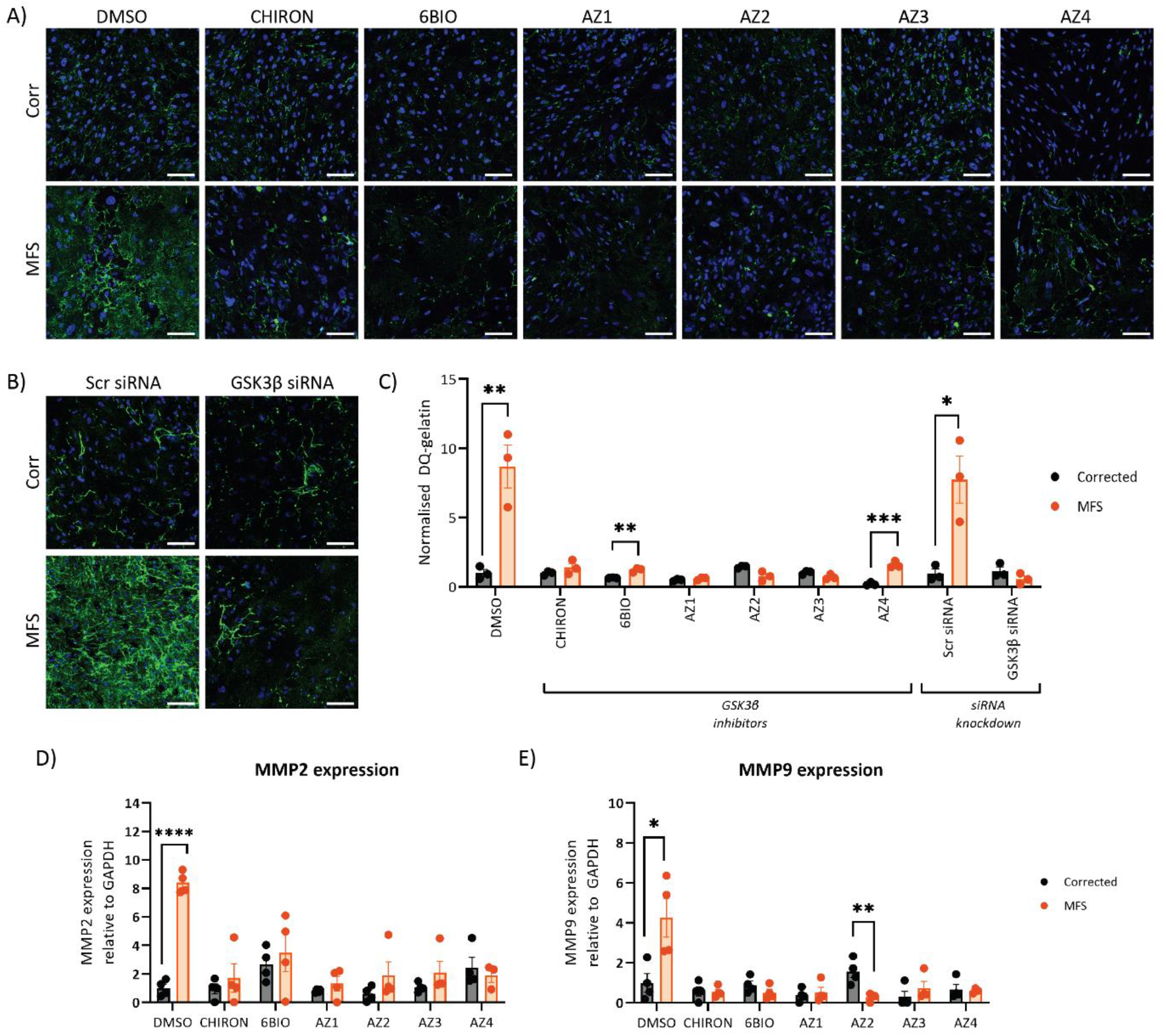
Decreased proteolysis and MMP expression upon disruption of GSK3β. *In-situ* gelatin degradation assay with either treatment of GSK3β SMI 1μM **(A)** or siRNA **(B)**, and quantification **(C)** n=3. Analysis of mRNA expression of MMPs 2 and 9 **(D)** and **(E)** indicate that their expression is decreased following SMI treatment; n=3-4. 150μm scale bars throughout. Data are represented as mean ± SEM. Cells treated under control condition (DMSO) was also used as controls for Supplementary Figure 3.

### GSK3β inhibition reduces apoptosis

Next, we aimed to see whether GSK3β inhibition could also decrease apoptosis using TUNEL staining. We confirmed that the assay was working by treating cells with DNase I (Supp Figure 4A), and non-GSK3β SMI (Supp Figure 4B). As before, after treatment for 96 hours with SMI (1μM) or siRNA against GSK3β, we fixed and stained cells (Figure 5). MFS VSMCs with control treatments had a higher percentage of apoptotic cells compared to the corrected control. Treatment with SMIs and siRNA improved this disease phenotype, with the exception of SMI AZ4 (Figure 5).

**Figure 5.**
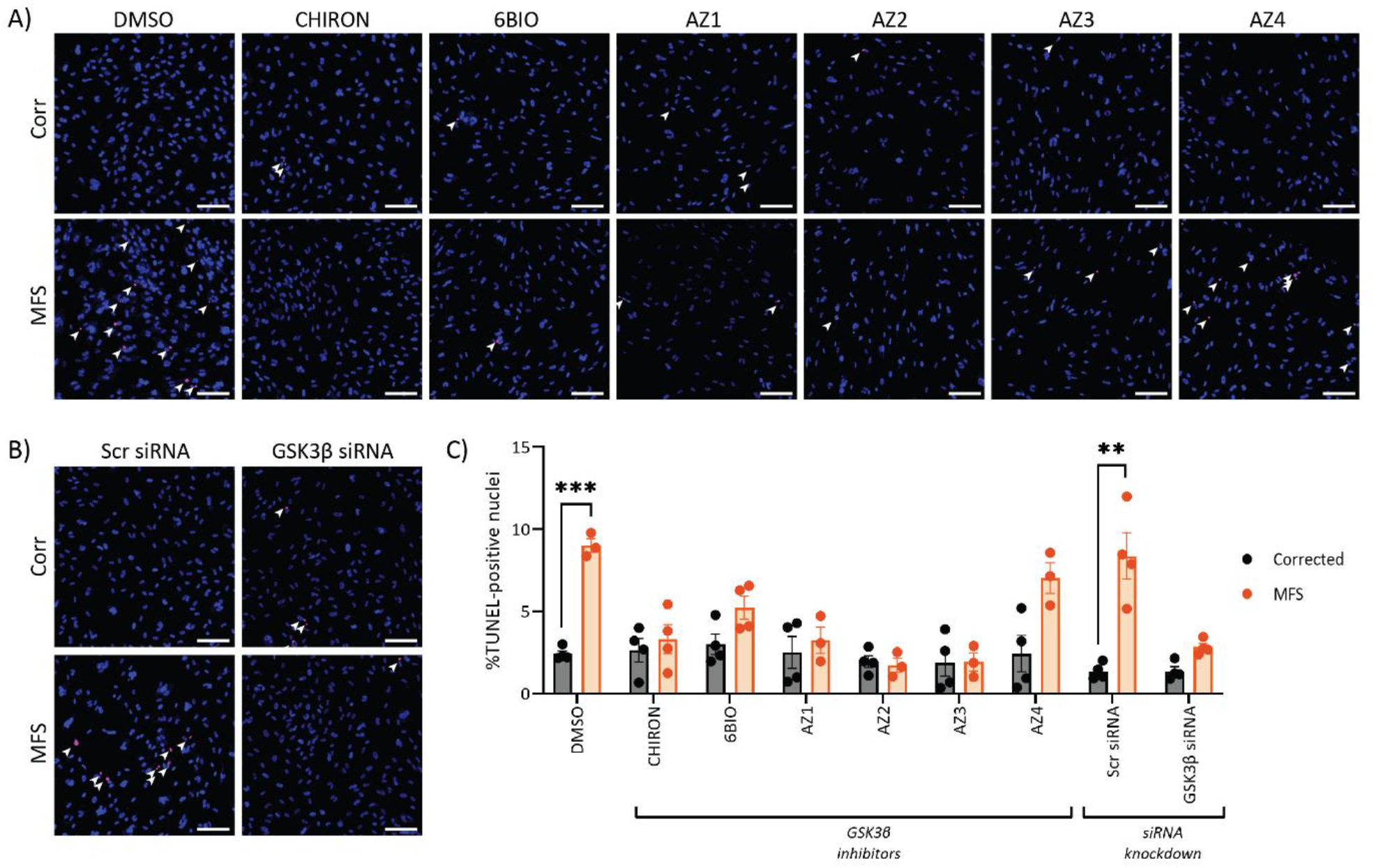
GSK3β inhibition or knockdown decreases apoptosis in MFS VSMCs. **A & B)** Cells were treated for 96 hours prior to staining for TUNEL (red) and DAPI (blue). **C)** Quantification was performed in a blinded and unbiased way using ImageJ and a macro. 150μm scale bars throughout. n=3-4. Data are represented as mean ± SEM. Cells treated under control condition (DMSO) was also used as controls for Supplementary Figure 4.

### Proliferation is unaffected by GSK3β

We subsequently aimed to determine whether GSK3β inhibition could improve the proliferation phenotype by performing EdU incorporation analysis. We cultured cells in the presence of EdU for 16 hours on the last day of SMI or siRNA treatment. Since the VSMCs we produce are not highly proliferative, we used HS27a cells as a positive control, and confirmed that our EdU signal coincided with KI67 staining (Supp Figure 5A). We noted that without treatment, MFS VSMCs had very poor proliferation, consistent with our experience when culturing them. In contrast, isogenic control cells had approximately 10% of cells synthesising new DNA. Unfortunately, neither treatment with GSK3β SMI nor GSK3β siRNA rescued the proliferation defects (Supp Figure 5).

### GSK3β inhibition reduces proteolysis and apoptosis in three additional MFS patient lines

Finally, we also sought to determine whether GSK3β inhibition is also beneficial in additional MFS patient iPSC lines. Three additional lines were used for this validation: DE35, DE37 and DE119 (Supp 1). These patients were diagnosed with MFS and experienced an aortic event, either dissection/rupture or surgery to replace a part of the aorta, before 18 years of age. These patient lines were reprogrammed into iPSCs, differentiated into NC-VSMCs and treated with GSK3β-targeting SMI AZ3 at 1μM as was done previously. AZ3 was selected over the other compounds, as we noted it had the fewest off-target effects (Supp Figure 6). Cell phenotype was assessed by looking at DQ-gelatin fluorescence and %TUNEL-positive nuclei (Figure 6).

**Figure 6.**
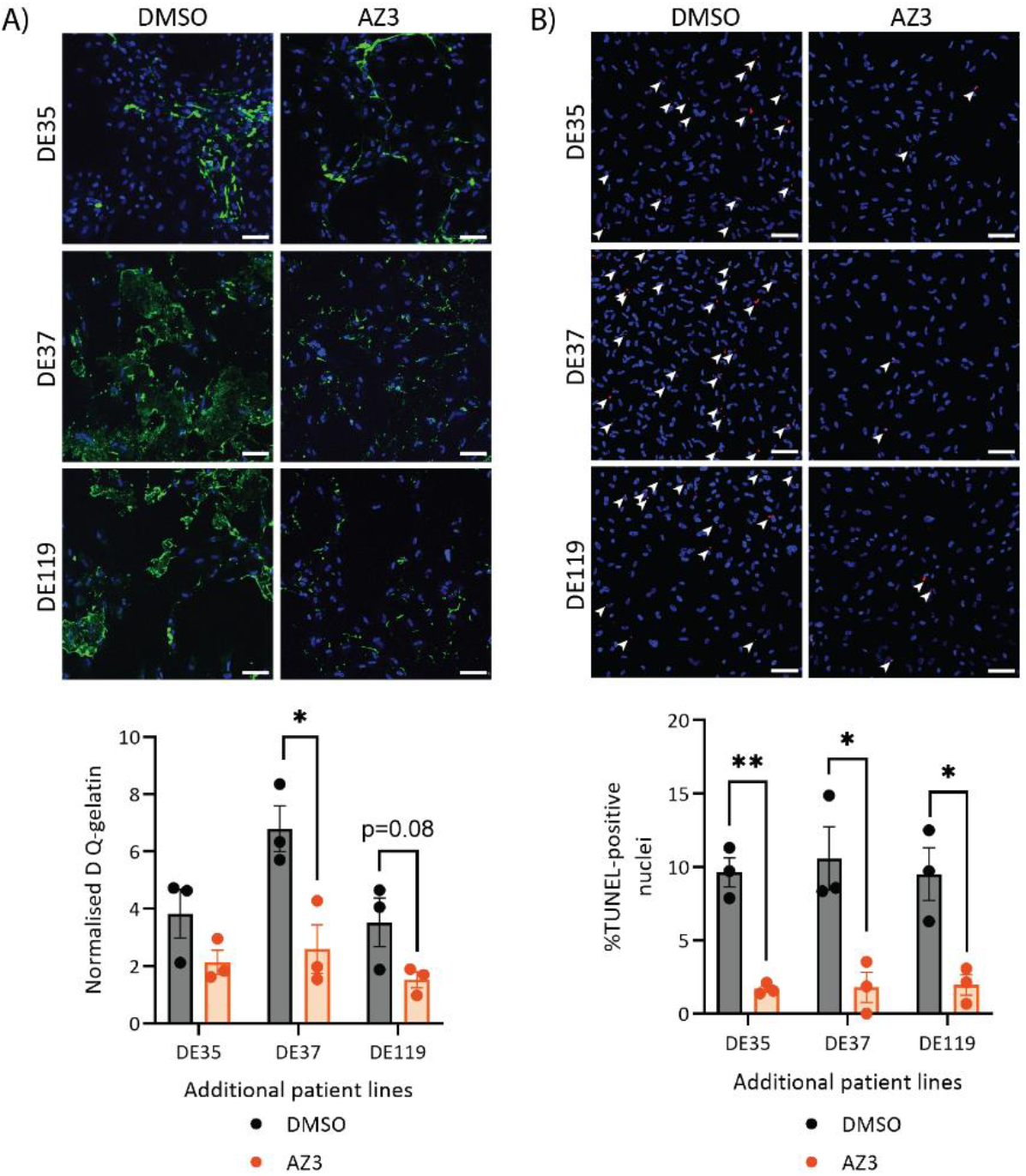
Inhibition of GSK3β using SMI AZ3 in three additional MFS patient lines is beneficial. Three additional patient lines – DE35, DE37 and DE119 – were differentiated into NC-VSMCs and treated with AZ3. Assays were performed as before to assess the effect of GSK3β inhibition on **A)** proteolysis and **B)** apoptosis. 150μm scale bars throughout. Data are represented as mean ± SEM.

The results with these three additional lines support what we have demonstrated with the C1242Y line. In terms of MMP activity, while the decrease in DQ-gelatin signal was only statistically significant in DE37, we noted that there was a trend towards decreased proteolysis in the two other lines with AZ3 treatment (Figure 6A). Equally, apoptosis was dramatically decreased with GSK3β inhibition (Figure 6B). Taken together, this work suggests that GSK3β could be a valuable target to further pursue.

## Discussion

### GSK3β activity in aortic aneurysms

The role of GSK3β in the development of aortic aneurysms is not entirely clear. There is evidence of increased GSK3β phosphorylation in abdominal aortic aneurysms (AAA) (Krishna et al., 2017). GSK3β was also identified as a likely regulator of pathogenic mechanisms from analysis of perivascular adipose tissue of patients with AAA (Piacentini et al., 2020). Here, we demonstrated that GSK3β inhibition was beneficial in our iPSC-derived MFS VSMCs using both multiple SMIs and a genetic approach. In addition, we have validated that SMI inhibition of GSK3β is beneficial in three additional MFS patient lines, suggesting that this may be a common disease mechanism and not a defect specific to the cell line that we used for the initial screening.

GSK3β activity is regulated in an unconventional way compared to most kinases. Many of its targets need to be primed with phosphorylation by another kinase; this post-translational modification will then fit within a groove of GSK3β, allowing it to phosphorylate its target. Inhibitory phosphorylation of GSK3β at the N-terminal Ser9 results in an autologous pseudo-substrate, preventing its binding to primed substrates (Frame et al., 2001). In this study, we have identified that the expression of GSK3β and its phosphorylation at Ser9 is paradoxical, with MFS VSMCs expressing less total GSK3β and having more of the inhibitory Ser9 post-translational modification when compared to the isogenic corrected control. Therefore, a big question is why GSK3β inhibition was beneficial, despite there being less total GSK3β and more inactivated GSK3β in MFS VSMCs.

There may be explanations that account for this paradox. First, Ser9 phosphorylation may not be a direct readout for GSK3β activity. As reviewed thoroughly by the Jope group, there are four main reasons why this may be: 1) not all GSK3β substrates are primed, 2) GSK3β is often found in complex with other proteins and p-Ser9 does not affect its activity within protein complexes, 3) p-Ser9 does not cause total inactivation of its activity and 4) its subcellular localisation could also impact how p-Ser9 affects activity (Beurel et al., 2015). In addition, the observed levels of GSK3β could also be the cells’ attempt to incompletely compensate for abnormal cell signalling; treatment with SMIs or siRNA would decrease the need for such compensation, thereby reducing some of the disease phenotypes.

### Downstream targets of GSK3β and clinical perspective

GSK3β is a kinase with numerous interacting partners. While most kinases have an average of 12 interacting partners, GSK3β is predicted to have over 500 targets (Linding et al., 2009). This is due to the unique mechanisms which regulate the activity and availability of GSK3β in a given cell. In this study, we have demonstrated that inhibition of this multi-target kinase with multiple compounds is beneficial, although GSK3β is not a straightforward enzyme to target in a clinical environment. It is ubiquitous and highly expressed in a number of organs, leading to concerns over toxicity and chronic usage. Yet lithium salts, which include GSK3β as one of their targets (Stambolic et al., 1996) have been used for decades to treat psychiatric disorders (Freland and Beaulieu, 2012), demonstrating the feasibility of long term GSK3β inhibition at appropriate doses. The therapeutic window of lithium is quite narrow, between 0.4nmol/L and 0.8nmol/L, and doses above this threshold are not well-tolerated (Malhi and Berk, 2012). AZD1080, which was also used in our *in vitro* studies under the name “AZ3” (Table 1), has progressed into Phase I clinical trials (Georgievska et al., 2013), although was subsequently abandoned after finding that it resulted in abnormalities in dog gall bladder at that dosage and did not enter Phase II trials (Bhat et al., 2018). Currently, one GSK3β inhibitor, Tideglusib, has successfully gone through Phase II trials for myotonic dystrophy (Horrigan et al., 2020).

The question of toxicity is particularly important for treating a life-long disease such as MFS. Upon diagnosis with the disease, patients will likely continue drug treatments for the rest of their lives. As such, treatment regimens have to be extremely well-tolerated. Losartan was unsuccessful in clinical trials, yet the very similar drug irbesartan retarded the rate of aortic growth compared to the placebo (Mullen et al., 2020). One key difference between losartan and irbesartan is their respective half-lives: the longer half-life of irbesartan increases its bio-availability compared to losartan, suggesting that insufficient dosage could be one of the reasons behind poor performance of losartan in clinical trials. With this in mind, a GSK3β inhibitor alone may not be appropriate for treating MFS – instead, combining GSK3β inhibitors with other drugs, all at lower concentrations, would allow us to target multiple signalling abnormalities whilst still being well-tolerated by patients. Alternatively, there are likely a multitude of downstream effectors which are currently unknown but may be more specific in the context of MFS and other aortic diseases. Proteomics and other unbiased approaches could be used to identify downstream effectors of the GSK3β pathway in the aorta which may be more tractable to therapeutic intervention.

### Further optimising the phenotypic screen

Despite recent demonstration of the benefits of angiotensin-II receptor blockers, there is still enormous scope for additional therapeutic intervention since irbesartan did not reverse or halt progressive aortic dilatation. Moreover, the experience with losartan suggests that mouse models can be difficult to align with the results of clinical trials. As a result, screening for potential therapeutic compounds solely in mice is not efficient.

Using this human *in vitro* screen, we have identified GSK3β among other interesting targets. The screen in this assay was performed in a medium-throughput manner where cells were cultured in a 24-well format. Scaling cell culture to 96-well plates for higher throughput was unsuccessful for the initial drug screen; we hypothesise that this is an effect of cell density and insufficient signal-to-noise ratio. We have since managed to perform MMP activity assays in a 96 well format by modifying the culture conditions. This strategy could therefore be used for future studies allowing us to interrogate even larger libraries of compounds, obtain a larger list of potential targets and generally improve the power of our compound screens.

Our screen has identified a number of compounds which were able to decrease the proteolytic phenotype of MFS. However, even from our relatively modest drug screen of 1,022 compounds, we identified 538 unique putative drug targets where pIC50 values were above 6. In future studies, we aim to interrogate a much larger cohort of SMIs, which in turn will also generate a larger list of potentially drug-able targets. How this large list is managed and how one decides which compounds are worthwhile to pursue will be a challenge. One strategy could be to compare the list of interesting drug targets to lists of SNPs which have been identified in genome-wide studies. Another strategy could be to follow up the compound screen with a genetic screen, utilising CRISPR for example, to narrow down the list of putative targets even further.

When further optimised, we envisage that this screening strategy could be an extremely valuable toolkit for future studies. As discussed above, a combinatorial approach to treating disease could be explored, where patient-derived VSMCs are treated with different combinations of drugs, all at doses chosen to minimise *in vivo* toxicity. Additional patient lines could also be studied to identify signalling pathways which are commonly disrupted – these particular pathways could potentially be very interesting when considering new clinical trials. Finally, it could be expanded towards other aortic diseases which exhibit abnormalities in MMP activity, such as Loeys-Dietz syndrome (Sinha lab, unpublished data).

### Experimental procedures

#### Cell culture

Isogenic control and patient iPSC lines were derived, cultured and differentiated as described previously (Cheung et al., 2012; Granata et al., 2017; Serrano et al., 2019). The patient line contains a C1242Y mutation in FBN1, and the original fibroblast line was obtained from Coriell cell bank (GM21943), and the isogenic control was generated using CRISPR-Cas9. Additional patient lines (denoted DE35, DE37 and DE119) were obtained from Sonalee Laboratory, St George’s Hospital, London, UK with the help of Dr Anne Child. These were received as fibroblasts and were reprogrammed using Sendai Virus V2.0, as performed previously (Granata et al., 2017). Briefly, iPSCs were cultured and maintained on vitronectin-XF (Stem Cell Technologies) and E8 media [DMEM/F12 (Gibco), Insulin-Transferrin-Selenium Supplement (Gibco), 0.44μM L-ascorbic acid (Sigma), 0.05% sodium bicarbonate (Sigma-Aldrich), 25ng/ml FGF2 (R&D Systems) and 1.74ng/ml TGF-β (Peprotech)]. For differentiation, a chemically-defined medium (CDM) [50% IMDM (Gibco), 50% Ham’s F12 Nutrient Mix (Gibco), chemically-defined lipid concentrate (Life Technologies), 15μg/ml transferrin (R&D Systems), 7μg/ml insulin (Sigma-Aldrich), 450μM monothioglycerol (Sigma-Aldrich) and 1mg/ml poly-vinyl alcohol (Sigma-Aldrich)] was supplemented with different cytokine and inhibitors. NC differentiation was initiated by culturing iPSC colonies in FSB media [CDM with 12ng/ml FGF2 (R&D Systems) and 10nM SB431542 (R&D Systems)] for 4 days, before being split into single cells and further cultured on 0.1% gelatin-coated plates. These NC were cultured and differentiated into NC-VSMCs in PT media [CDM with 10ng/ml PDGF-BB (Peprotech) and 2ng/ml TGF-β (Peprotech)] for 12 days. After differentiation, VSMCs were matured for 2 weeks in DMEM/F12 (Gibco) containing 10% foetal bovine serum (Gibco) before being used in assays (Figure 1A).

#### SM screen

SMs were obtained from AstraZeneca. 1,022 drugs were selected out of their library of 14,000 compounds (Parafati et al., 2020). These SMs were diluted from 10mM stock in DMSO to a final concentration of 1μM in MEF media. Control and MFS NC-VSMCs were seeded onto 0.1% gelatin-coated 24-well plates. The following day, the 96 hour treatment with SMIs began, with a medium refresh halfway through. On day 4, cell culture medium was then collected to assay for MMP activity using the SensoLyte® 520 Generic MMP Assay Kit Fluorometric (Anaspec) according to the manufacturer’s instructions for protocol B. Briefly, supernatants were incubated with 1mM AMPA for 3 hours at 37°C and 50μl were transferred to a 96-well plate. 50μl of the included MMP substrate solution was added to each well and further incubated for 1 hour at room temperature after which 50μl Stop Solution was added to terminate the reaction. Fluorescence intensity, corresponding to MMP activity, was measured at Ex/Em = 490nm/520nm on a plate-reader.

#### siRNA transfection

siRNA knockdown was performed in Opti-MEM media (Gibco) and Dharmafect 1 Transfection Reagent (Horizon Discovery). A non-specific siRNA (ON-TARGETplus; Horizon Discovery) was used as a control alongside siRNA against GSK3-β (Invitrogen). Knockdown in wells of a 6-well plate were performed by incubating 40nM siRNA with Dharmafect 1 for 20 minutes before applying to cells. The next day, cell culture media was refreshed, and cells grown for another 3 days before downstream experiments.

#### Immunocytochemistry (ICC)

Cells were washed with PBS and fixed with 4% PFA for 15 minutes. After subsequent washes with PBS, cells were permeabilised with 0.05% triton X-100 before blocking with 10% FBS in PBS. The primary antibody (Supp Table 2) was incubated overnight at 4°C. The next day, samples were washed in PBS and incubated with the secondary antibody (Supp Table 1) for 1 hour at room temperature. Finally, after another wash in PBS, nuclear staining was performed using Hoechst 33342 (Invitrogen) for 10 minutes.

#### DQ-gelatin assay

DQ-gelatin fluorescein conjugate (Invitrogen) was dissolved in water to 0.5mg/ml and used to coat Ibidi 8-well chambered slides or 96-well plates for 24 hours at 4°C. Dishes were washed twice with PBS before seeding 15,000 VSMCs. The following day, cells were treated with either SMIs or transfected with siRNA for 96 hours before washing with PBS and fixing in 4% PFA (Alfa Aesar) for 10 minutes at room temperature. Fixed cells were subsequently imaged using a Zeiss LSM 710 confocal microscope. Resulting images were processed and quantified in ImageJ. DQ-gelatin fluorescence intensity was determined after image processing and thresholding. The number of nuclei was also determined after initial processing and analysis of particles. All described image processing and quantification steps were performed using a macro for automated and unbiased analysis.

#### TUNEL staining

VSMCs were seeded onto 0.1% gelatin-coated plates and were treated with either SMIs or transfected with siRNA for 96 hours. TUNEL staining to identify apoptotic cells was performed using the In Situ Cell Death Detection Kit (Roche) according to the manufacturer’s instructions. Positive controls were obtained by treating cells with 3U/ml DNase I (Sigma-Aldrich). Tiled images were taken using a Zeiss LSM 710 confocal microscope and quantified in ImageJ using a macro. After image processing, the number of TUNEL-positive nuclei was quantified.

#### EdU proliferation assay

VSMCs were seeded onto 0.1% gelatin-coated plates and began their 96-hour treatment with either SMIs or transfection with siRNA. They were then incubated with 20μM EdU for 16 hours. EdU staining was performed using the Click-iT™ Plus EdU Cell Proliferation Kit (Invitrogen) as per the manufacturer’s instructions. Images were taken using an automated Leica Matrix DMI 6000 microscope. Images were processed and quantified in ImageJ. The number of nuclei stained either for EdU or Hoescht was quantified after initial image processing. All steps were performed using a macro.

#### RNA extraction and RT-qPCR

After washing the cells with PBS, RNA extraction was performed from cells growing in 12-well plates using the GenElute Mammalian Total RNA Miniprep Kit (Sigma-Aldrich) according to the manufacturer’s instructions for extraction from adherent cells. After quantification, reverse transcription was performed using the Maxima First Strand cDNA Synthesis Kit (Thermo Scientific). RT-qPCR was performed using SYBR Green (Applied Biosystems) with 5ng cDNA per sample. Experiments were performed with technical triplicates, and gene expression was determined based on the expression of housekeeping gene GAPDH using the ΔCT quantification method.

#### Protein extraction and western blotting

Protein was extracted from adherent cells with RIPA buffer (Sigma-Aldrich) supplemented with HALT Protease and Phosphatase Inhibitor (Pierce). After quantification using the BCA Assay (Pierce), lysates were then mixed with 4x Sample Buffer (Bio-Rad) containing β-mercaptoethanol before being denatured at 98°C for 5 minutes. Lysates were subsequently centrifuged at 16,000 x g for 5 minutes, and supernatant corresponding to 4μg protein was then loaded into 4–15% Mini-PROTEAN® TGX™ Precast Protein Gels (Bio-Rad). SDS-PAGE was run for approximately 90 minutes at 80V. Proteins were transferred on a methanol-activated PVDF membrane (Merck) using the wet transfer method running at 60V for 2 hours. Membranes were rinsed with Tris-buffered saline plus 0.1% Tween 20 (TBST) before blocking in 5% BSA for 1 hour before primary antibody incubation was performed overnight at 4°C (Supp Table 3). The following day, membranes were washed 3 times with TBST for 10 minutes each and incubated with HRP-conjugated secondary antibodies (Supp Table 2) for 1 hour at room temperature. Membranes were washed another 3 times with TBST for 10 minutes each before incubating with ECL reagent (Pierce). Films were exposed to membranes before being developed using the Konica Minolta SRX-101A.

#### Drug target analysis

Drug target analysis was done using R (version 4.0.5) and the following packages: ggplot2, pheatmap, dplyr, tidyr, biomaRt and clusterProfiler (Durinck et al., 2009; Wickham, 2011; Wickham et al., 2019; Wu et al., 2021; Yu et al., 2012).

#### Statistics

A biological replicate is defined as VSMCs derived from distinct and separate NC differentiations. VSMCs obtained from a single NC differentiation are considered to be technical replicates. Statistical significance was determined using an unpaired two-tailed Student’s t-test, with p-values < 0.05 considered to be significant.

## Acknowledgements

This work was supported by the following British Heart Foundation grants: BHF Program grant (RG/17/5/32936) for HD; BHF Centre of Regenerative Medicine (RM/13/3/30159) for MM; BHF PhD Studentship (TG/18/4/33770) for RGCM and BHF Senior Fellowship (FS/18/46/33663) for SS.

We thank Dr Peter J. Holt for his helpful comments on the manuscript.

We also thank members of the iPSC core facility at the Wellcome-MRC Stem Cell Institute for their work reprogramming cell lines DE35, DE37 and DE119. This core facility was supported by the NIHR

Cambridge Biomedical Research Centre (BRC-1215-20014*). The views expressed in this manuscript are those of the authors and not necessarily those of the NIHR or the Department of Health and Social Care.

JAAM and AC thank the Marfan syndrome patients who participated in this study, and the Marfan Trust for funding their work.

The authors gratefully acknowledge the Advanced Microscope Facility, JCBC, for their support & assistance in this work, and in particular Darran Clements for his guidance.

This research was funded in whole, or in part by Wellcome Trust (203151/Z/16/Z) and the UKRI Medical Research Council (MC_PC_17230). For the purpose of open access, the author has applied a CC by public copyright licence to any Author Accepted Manuscript version arising from this submission.

## Author contributions

HD conceived, performed experiments and analysis, and wrote the paper. MM and RGCM performed the drug screen with supervision and guidance from AG. MH performed blotting experiments. MF and DMS assisted with the design of the screen, provided compounds and assisted with analysis. AC provided patient phenotypes and cell lines from classical Marfan syndrome patients, and JAAM provided fibrillin-1 mutations. S.S. conceived and supervised the project. All authors reviewed the manuscript.

## Declaration of interests

The authors declare that the research was conducted in the absence of any commercial or financial relationships that could be construed as a potential conflict of interest.

## Notes

### Competing Interest Statement

The authors have declared no competing interest.

