## Supplemental Figures and Tables for "A phenotypic screen of Marfan syndrome iPSC-derived vascular smooth muscle cells uncovers GSK3β as a new target"

### Supplemental Figures & Tables

A)

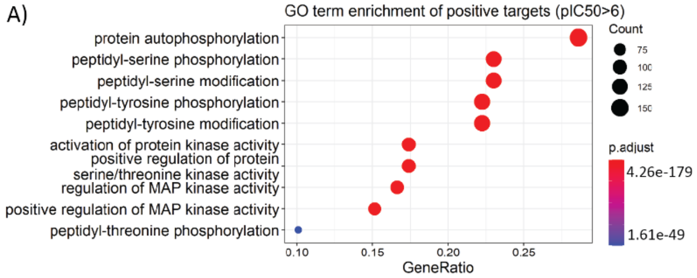


B)

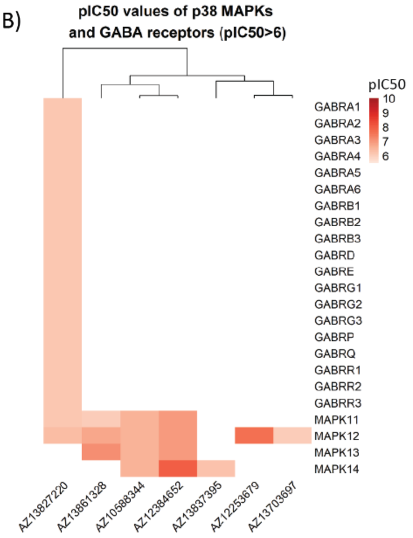


C)

| ID | Description | pvalue | p.adjust | qvalue | Count |
|---|---|---|---|---|---|
| hsa04010 | MAPK signaling pathway | 3.50E-39 | 8.72E-37 | 2.95E-37 | 82 |
| hsa04068 | FoxO signaling pathway | 5.03E-30 | 6.26E-28 | 2.12E-28 | 49 |
| hsa04012 | ErbB signaling pathway | 2.47E-29 | 2.05E-27 | 6.93E-28 | 40 |
| hsa04722 | Neurotrophin signaling pathway | 5.77E-29 | 3.59E-27 | 1.21E-27 | 46 |
| hsa05135 | Yersinia infection | 6.85E-23 | 3.41E-21 | 1.15E-21 | 43 |
| hsa05161 | Hepatitis B | 2.21E-22 | 9.17E-21 | 3.10E-21 | 46 |
| hsa04660 | T cell receptor signaling pathway | 5.65E-22 | 2.01E-20 | 6.79E-21 | 37 |
| hsa04360 | Axon guidance | 6.14E-21 | 1.91E-19 | 6.46E-20 | 47 |
| hsa04510 | Focal adhesion | 1.28E-20 | 3.53E-19 | 1.19E-19 | 49 |
| hsa05170 | HIV1 infection | 2.38E-20 | 5.93E-19 | 2.01E-19 | 50 |

D)

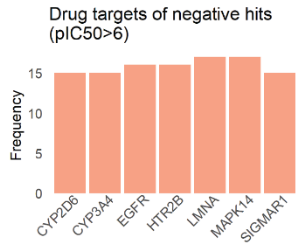


**Supplemental Figure 1. Computational analysis of positive hits from drug screen. A)** GO term enrichment of drug targets indicates that the majority of hits are for kinases. **B)** Previously identified p38 MAPK and GABA receptors were also found to be positive hits from the drug screen. Heatmap illustrates pIC50 values for these interactions. **C)** KEGG Pathway enrichment analysis. **D)** Top recurring targets of the negative hits from the screen does not include GSK3β.

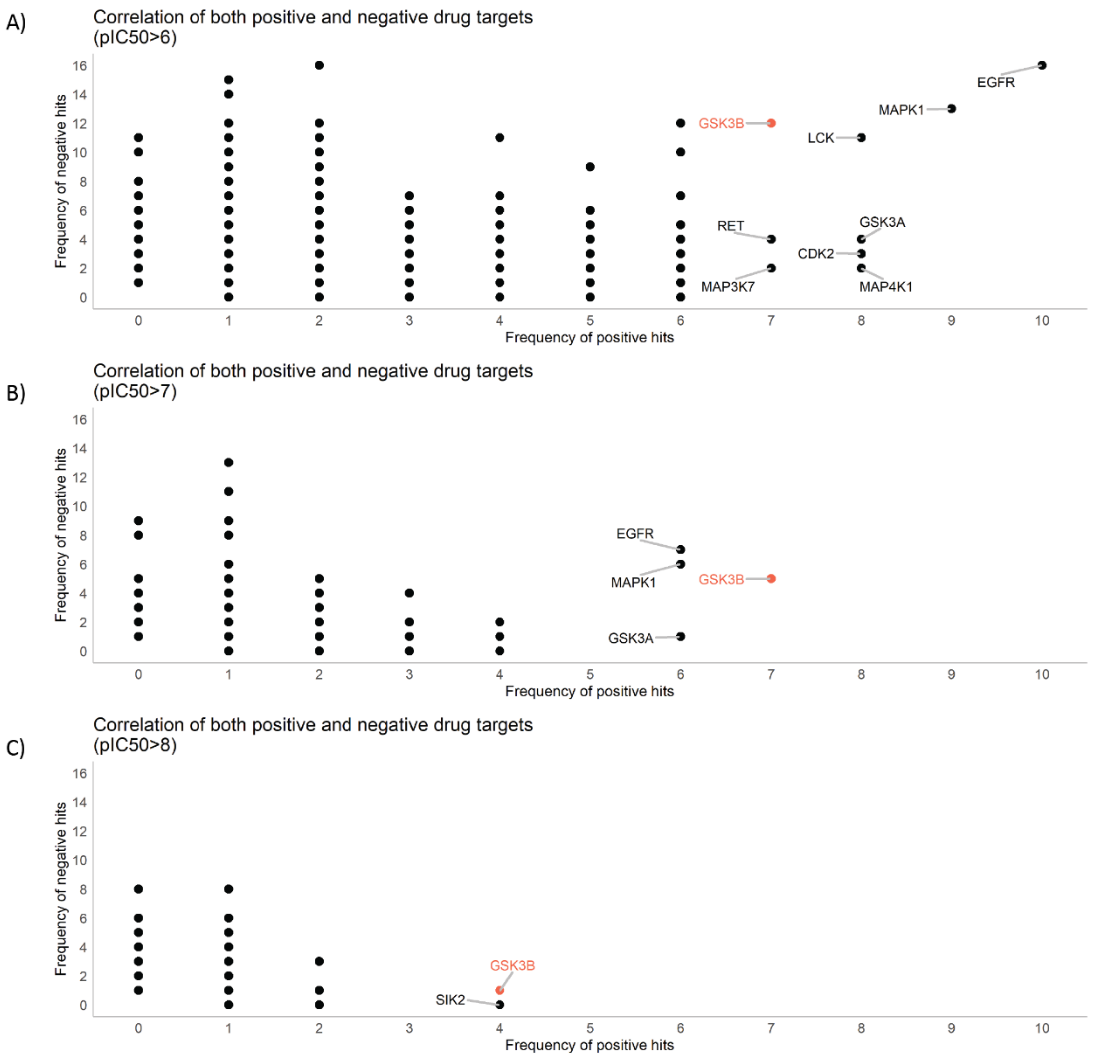


**Supplemental Figure 2. Correlation between the number of positive and negative hits filtered by pIC50>6 (A), pIC50>7 (B) and pIC50>8 (C).**

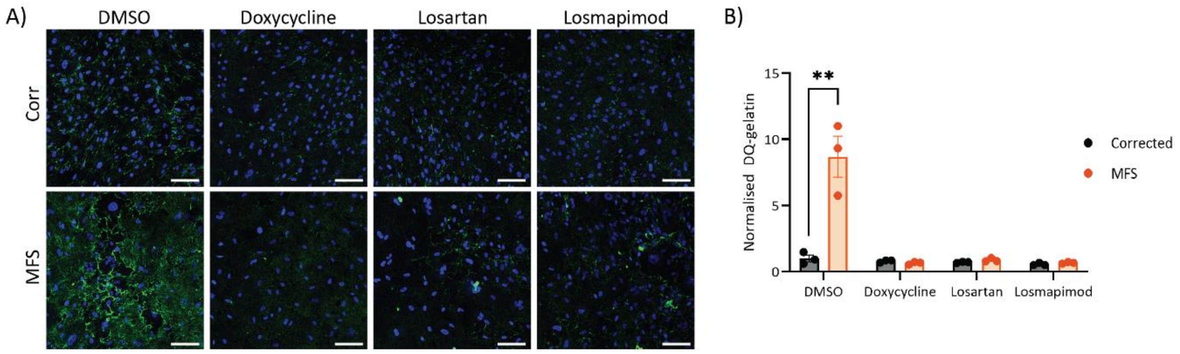


**Supplemental Figure 3. In-situ zymography with non-GSK3β inhibitors.** DQ-gelatin staining **(A)** and quantification **(B)**. 150µm scale bars throughout. n=3. Cells treated under control conditions (DMSO) were also used as controls for Figure 4.

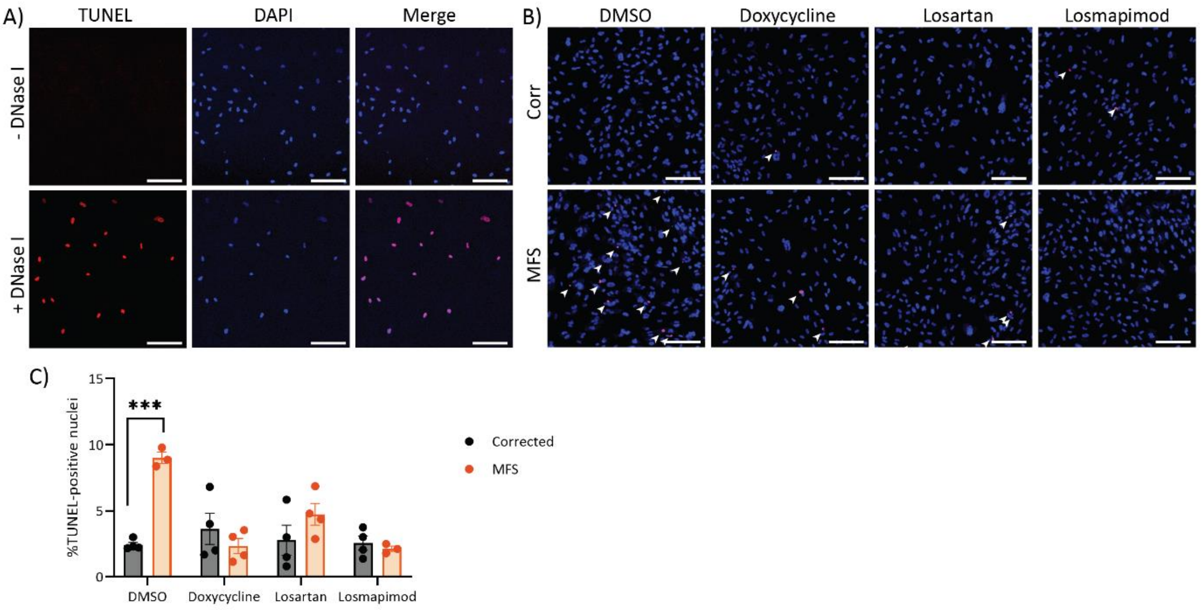


**Supplemental Figure 4. TUNEL staining to determine apoptosis. A)** Corrected VSMCs treated with DNase I show TUNEL staining. **B)** Treatment with non-GSK3β inhibitors and quantification of TUNEL% nuclei **(C)**. 150μm scale bars throughout. n=3-4. Cells treated under control condition (DMSO) was also used as controls for Figure 5.

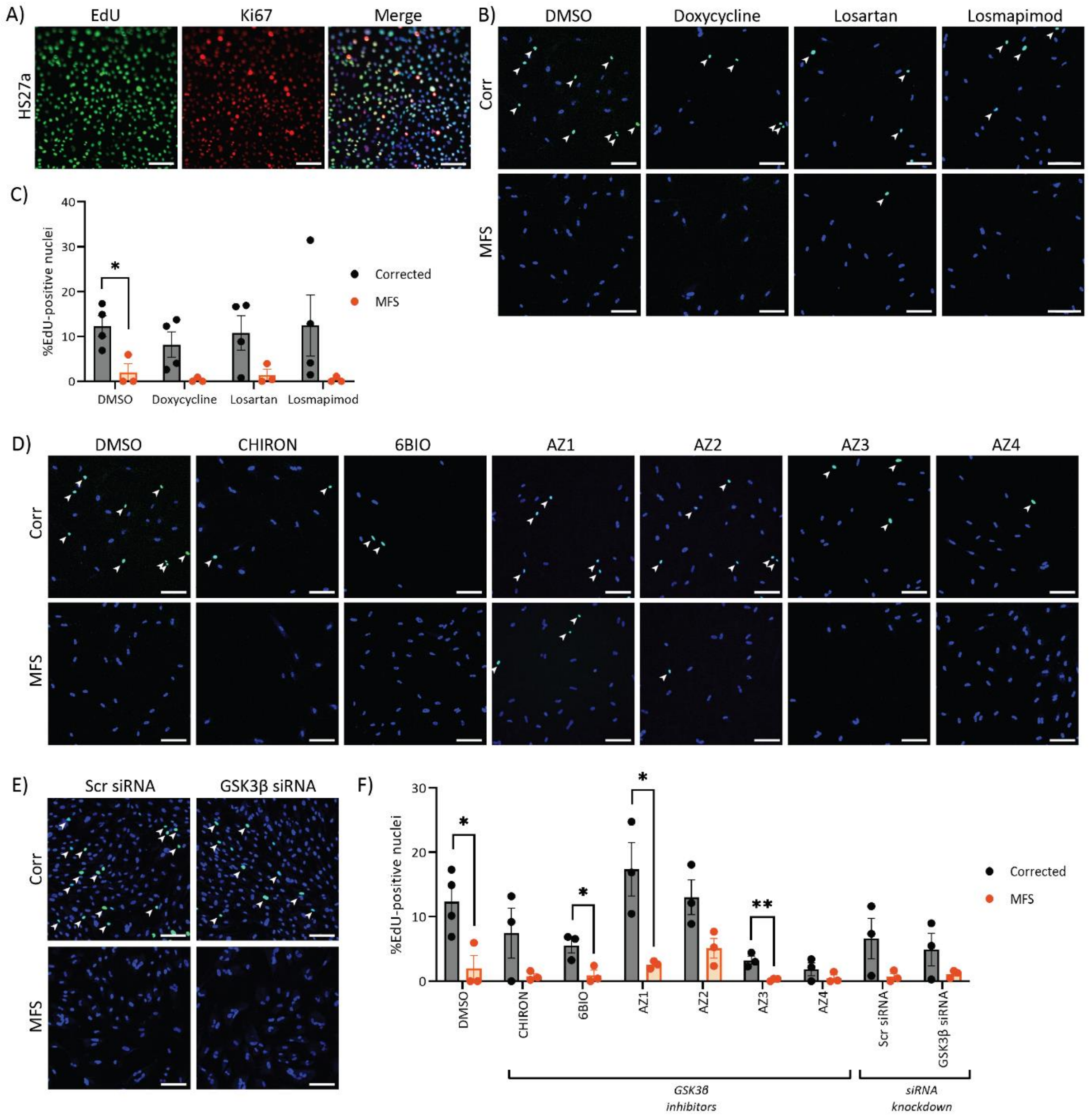


**Supplemental Figure 5. Proliferation was assessed by incorporation of EdU. A)** Highly-proliferative HS27a cells were used as a positive control was used to verify staining conditions. Incorporated EdU was stained in green, Ki67 in red and DAPI in blue. **B)** Effect of non-GSK3β inhibitors on proliferation and **C)** quantification. **D)** Effect of GSK3β inhibitors and siRNA on proliferation and **F)** quantification. n=3-4. 150µm scale bars throughout. Cells treated under control condition (DMSO) were used as controls for panels B and D and their quantifications.

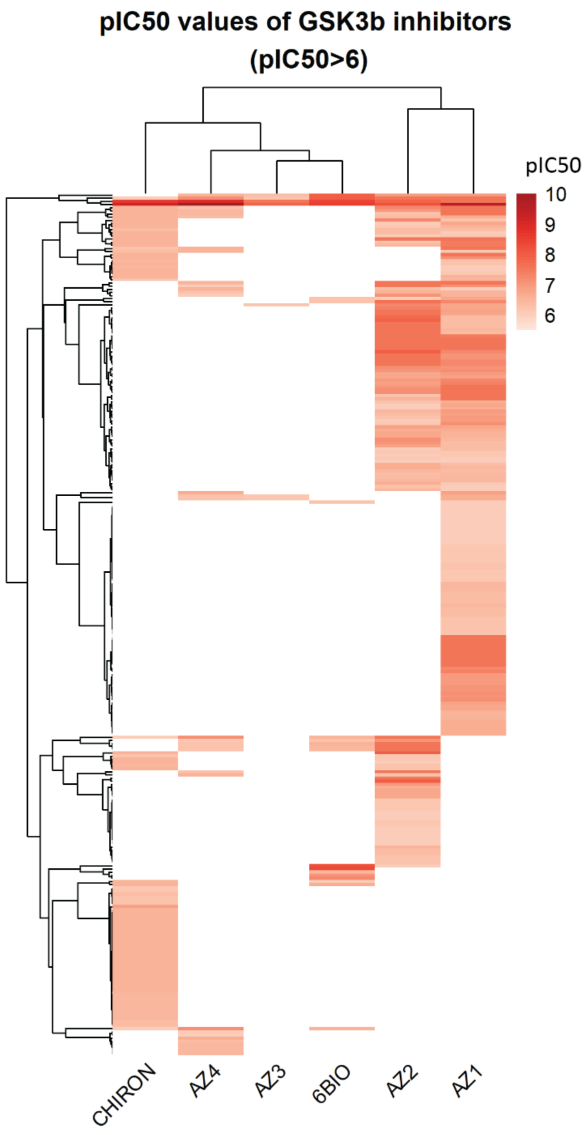


**Supplemental Figure 6. Off-targets of GSK3β SMI and their pIC50 values.** pIC50 values are represented in a heatmap.

| **Cell line** | ***FBN1* mutation** |
|---|---|
| DE35 | c.1837+5G>C |
| DE37 | Unknown – diagnosis was based on clinical criteria. Reduced fibrillin-1 deposition in patient fibroblasts. |
| DE119 | c.1051C>T; p.(Q351*) |

**Supplemental Table 1.** Additional MFS patient lines used. Where *FBN1* mutations were unavailable, diagnosis was based on clinical criteria for MFS.

| **Antibody target** | **Catalogue number** | **Supplier** | **Concentration** |
|---|---|---|---|
| CNN1 | C2687 | Sigma | 1:300 |

| | | | |
|---|---|---|---|
| KI67 | 9129 | Cell Signalling Technology | 1:300 |
| Goat anti-mouse IgG (Alexa Fluor 594) | A32742 | Invitrogen | 1:1,000 |
| Goat anti-rabbit IgG (Alexa Fluor 594) | A32740 | Invitrogen | 1:1,000 |

**Supplemental Table 2.** Antibodies used for ICC.

| Antibody target | Catalogue number | Supplier | Concentration |
|---|---|---|---|
| GSK3β | 9832 | Cell Signalling Technology | 1:1,000 |
| GSK3β (Ser9) | 9336 | Cell Signalling Technology | 1:1,000 |
| GSK3α | 9338 | Cell Signalling Technology | 1:1,000 |
| β-catenin | 9562 | Cell Signalling Technology | 1:1,000 |
| GAPDH | G9545 | Sigma-Aldrich | 1:10,000 |
| Mouse secondary | NA931 | GE-Healthcare | 1:5,000 |
| Rabbit secondary | 7074 | Cell Signalling Technology | 1:2,000 |

**Supplemental Table 3.** Antibodies used for western blotting.

#### Data

Data 1. pIC50 values of drug targets from positive hits from the drug screen.

Data 2. Frequency of drug targtes amongst effective SMs from drug screen.

Data 3. pIC50 values of drug targets of GSK3β-targetting SMIs used for validation.
